# MetaTree: an interactive web platform for aligned hierarchical data visualization and multi-group comparison

**DOI:** 10.64898/2026.01.22.701099

**Authors:** Qing Wu, Ailing Zhang, Zhibin Ning, Daniel Figeys

## Abstract

**Background:** Hierarchical quantitative profiles are widely used in microbiome studies and other domains. However, comparing multiple samples and experimental groups while preserving hierarchical structure remains challenging. Many existing workflows require extensive manual figure assembly or do not support aligned comparisons across conditions on a shared hierarchy.

**Results:** We developed MetaTree, an open-source platform that runs in a web browser for interactive visualization and comparative analysis of hierarchical quantitative data. MetaTree anchors samples, groups, and contrasts between groups to a shared reference hierarchy, preserving one-to-one node correspondence so that the same clade is compared in the same position across views. In addition to visualization, MetaTree integrates statistical testing for comparisons between two groups with false discovery rate (FDR) control, enabling users to identify clades with consistent differences between conditions and interpret them in hierarchical context. MetaTree also provides user configurable controls for visual encoding, filtering thresholds, label density, and layout, allowing figures to be adapted to different datasets and reporting needs. The interface remains usable for large hierarchies through interactive navigation, adaptive label handling, and branch collapsing.

**Conclusions:** MetaTree is an installation-free web platform (https://byemaxx.github.io/MetaTree) for topology-consistent visualization and comparison of hierarchical profiles, supporting coordinated multi-panel exploration and automated comparison matrices to enable rapid generation of publication-ready figures for microbiome and other hierarchical datasets.

## Background

Many biological datasets, including taxonomic hierarchies and functional classification systems [1-3], are organized across multiple levels. In microbiome workflows such as 16S rRNA sequencing [4], shotgun metagenomics [5] and metaproteomics [6], thousands of features are mapped onto these hierarchies. Interpreting how clades respond to perturbations therefore benefits from visualization methods that preserve hierarchical structure [7]. However, common approaches have significant limitations. Stacked bar charts typically require aggregation to coarse ranks and are constrained by the number of distinguishable colors, obscuring patterns at deeper and often biologically informative levels [8, 9]. Clustered heatmaps, even with dendrograms, separate the hierarchy from the quantitative matrix, requiring users to mentally reconcile the two and becoming difficult to read as feature counts increase [10].

Several tools support phylogenetic tree visualization, including PhyD3 [11], GraPhlAn [12], ETE Toolkit [13], Krona [14], and iTOL [15], but they primarily focus on rendering individual trees rather than systematic comparison across many samples or groups. Many require scripting or configuration files, and iTOL does not natively provide a topology-consistent multi-panel workflow, where all panels are rendered on a shared reference topology with one-to-one node correspondence [15]. Heat-tree visualizations partially address this gap by embedding quantitative values within the tree. Metacoder [9] introduced heat-tree displays for microbial community analysis, but its dependence on the R environment, static ggplot2 output [16], and limited label control make it challenging to scale to large or complex datasets. In practice, labels frequently overlap or become unreadable at deeper levels, and assembling multi-panel, multi-group figures often requires substantial manual editing of static plots [10].

Recent microbiome-focused platforms have also expanded interactive data exploration and downstream analysis. For example, Mian [17] provides web-based exploration of microbiome abundance tables together with statistical and machine-learning functions, MicrobiomeAnalyst [18] offers a broad framework for statistical, functional, and integrative microbiome analysis, and Snowflake [19] presents microbiome abundance data using an alternative graph-based visualization strategy. However, these platforms are not primarily designed for topology-consistent comparison of multiple samples and group contrasts on a shared hierarchical reference. As a result, a practical gap remains for workflows centered on aligned hierarchical comparison and figure generation.

To address these limitations, we developed MetaTree, a browser-based platform for interactive heat-tree visualization and comparison of hierarchical quantitative data. MetaTree provides an integrated workflow for data upload, exploration, and export of publication quality figures, without requiring programming. Implemented in JavaScript with D3.js [20], MetaTree constructs a shared reference topology from hierarchical items so that samples and groups can be displayed with consistent node alignment across views. Beyond visualization, it includes built-in two-group comparisons with FDR control, supports operational taxon–function (OTF) structures [21], and provides dedicated multi-group comparison modes to examine clade-level patterns across samples and experimental conditions within a single interactive framework.

## Implementation

### Visualization framework and interaction

MetaTree is implemented as a single-page web application and runs entirely in the user’s browser. It first constructs a global reference tree by merging all hierarchical paths present in the input, ensuring that every panel shares an identical topology and differs only in the mapped quantitative attributes. This enables direct, side-by-side comparison of many samples or groups without re-rooting or re-ordering trees.

Quantitative information is encoded through multiple visual channels that can be adjusted interactively:

- **Tree color (nodes and branches)** encodes a user-selected variable such as abundance, intensity or log2 fold-change. MetaTree provides multiple built-in color palettes and allows users to adjust the value domain and the direction of the color scale. Transformations such as logarithmic or square-root scaling can be applied before mapping, making it possible to handle both highly skewed abundance distributions and centered contrast statistics.
- **Node size** reflects the magnitude of a quantity of interest, typically an aggregated abundance or intensity. Global size scaling controls allow users to emphasize or down-weight differences in node area for dense or sparse trees.
- **Branch width** can be used to enhance structural readability or to emphasize clades with higher overall signal, depending on the analysis goal.

MetaTree supports multiple layouts (radial, dendrogram, and circle packing) and provides panel-level controls for width/height, with optional synchronized resizing to facilitate figure assembly. Users can pan and zoom within panels; an optional synchronized zoom mode propagates navigation across panels, enabling aligned inspection of the same clade across many samples or contrasts.

To reduce clutter in deep hierarchies, MetaTree provides configurable label density (by depth and by abundance-driven thresholds) and a smart label-culling option that suppresses overlapping labels while preserving the underlying geometry. Targeted clade interrogation is supported through keyword search and an include/exclude filter list, as well as manual label-color overrides for highlighting specific taxa or functional categories.

For very large trees, readability and responsiveness are improved through interactive pruning. Users can collapse or expand branches by clicking internal nodes, reducing on-screen complexity while preserving global context; collapsed nodes are visually marked and can be restored via a dedicated control. Combined with include/exclude filtering and abundance-based thresholds, this enables progressive exploration from the full hierarchy to focused subtrees without changing the underlying reference topology.

Export is supported directly from the interface via per-panel context actions, producing publication-ready SVG (vector) or PNG (raster) outputs. Because all mapping parameters (palette, domain, transforms, label rules and filters) are applied interactively, exported figures reflect the exact visual state used during exploration.

### Data input and hierarchical model

MetaTree accepts two complementary table types to cover both raw hierarchical profiles and downstream statistical outputs:

- **Wide format**, rows represent hierarchical items and columns represent samples. Each row can be provided either as a delimited path (e.g. Kingdom; Phylum; Species) or as an explicit level-wise encoding, and MetaTree builds the tree by parsing the path into parent–child relationships.
- **Long format**, where each row encodes an item–sample or item–contrast pair with associated statistics, for example log2 fold-change, effect size and P-value obtained from statistical tools. This format enables direct visualization of statistical results (including significance-aware filtering) without requiring users to manually reshape outputs from external pipelines.

In addition to native wide and long table imports, MetaTree provides a converter workflow for commonly used external hierarchical formats. Supported converter inputs include BIOM [22] JSON, Newick trees with a sidecar abundance table, and QIIME [23] exported files. These inputs are converted into MetaTree-compatible tabular representations, which can be imported directly into MetaTree for one-click visualization or downloaded for downstream use.

MetaTree provides an interactive import wizard that maps user columns to the required semantic fields (hierarchy/path, sample or group labels, and value/statistic columns) and supports flexible delimiters and missing-value handling. Internally, items are keyed by their full ancestor path to avoid collisions when identical labels occur under different parents. Optional sample metadata can be loaded to define groups (e.g. treatment, time, cohort). Groups can also be defined and re-ordered interactively.

### Analysis modes

On top of the shared visualization framework, MetaTree implements four analysis modes:

- **Sample view (Individual samples):** multiple samples are shown simultaneously as separate heat-tree panels on the shared reference tree, with sample-specific values mapped directly to tree color and node size. This supports side-by-side inspection of sample-level heterogeneity and outlier detection.
- **Group view (Group samples):** samples are aggregated into user-defined groups based on metadata or interactive grouping, and group-level summaries (e.g. mean or median abundance) are visualized as one heat-tree per group. This mode facilitates comparison of overall hierarchical profiles across conditions.
- **Group comparison (Pairwise comparison):** two groups are selected for pairwise comparison. For each hierarchical item, MetaTree performs an exploratory between-group test using either the Wilcoxon rank-sum test [24] or Welch’s t-test [25], and controls the false discovery rate using the Benjamini-Hochberg (BH) procedure [26]. MetaTree reports log2 fold-change, defined as log2(Group 2 / Group 1) using group-level summary abundances, together with FDR-adjusted *P*-values. To ensure numerical stability when group summaries contain zeros, fold-changes are computed using an additive pseudocount. These statistics are used primarily for interactive filtering: users can restrict the display to items exceeding an absolute log2 fold-change threshold and passing an FDR cutoff, thereby focusing the tree on clades with consistent between-group differences.
- **Comparison matrix (All-versus-all):** when more than two groups are present, MetaTree can automatically compute pairwise comparisons between all group pairs and arrange the results in a matrix of heat-tree panels. Any cell can be selected for detailed inspection using the same visualization and filtering controls as in group comparison mode, providing an overview of relationships between conditions while preserving access to individual contrasts.

Across all modes, additional filters allow users to subset the tree by metadata (e.g. selected groups), by item identity (e.g. specific clades or keyword matches) and by simple thresholds on abundance or other numeric variables. P-values and FDR can be used as filtering criteria to control which items are shown, thereby reducing visual clutter and highlighting the most relevant features.

### Programmatic integration and deployment

Beyond the graphical interface, MetaTree exposes lightweight browser-side hooks for integration into analysis reports or dashboards. Data can be injected programmatically (e.g. via a JavaScript call that loads tabular text, or by embedding inline data in an HTML document), and a readiness event is emitted when the application has initialized. This design enables reproducible, shareable visualizations that remain fully functional when deployed as a static website (e.g. GitHub Pages) or run locally for offline, privacy-preserving analysis.

### Benchmark design

MetaTree performance was evaluated in a reproducible browser-based benchmark workflow implemented separately from the main application logic. Benchmarks were run in headless Chromium (Chrome 147.0.7727.101) on Windows x64 using Playwright automation on a system equipped with a 13th Gen Intel(R) Core (TM) i9-13900H processor, 32 GB RAM, and a 1600 × 1200 viewport. Synthetic benchmark datasets were derived from the example hierarchy to preserve realistic hierarchical structure and were generated at approximately 500, 2,000, 5,000, and 10,000 nodes. Across all benchmark settings, 72 conditions were evaluated with five repeated runs each (360 browser runs total). Measured quantities included data import time, hierarchy construction time, initial render time, filter-update time, collapse/expand update time, comparison computation time, comparison-matrix render time, and post-render JavaScript heap usage. Results are reported as medians and interquartile ranges.

## Results

### Interface overview and workflow

MetaTree provides a browser-based interface for interactive visualization and comparison of hierarchical quantitative data (Fig. 1). Users begin by loading either a native wide-format abundance table, optional sample metadata, or a long-format table of statistical results such as log2FC with significance metrics. To improve compatibility with common community formats, MetaTree also provides a converter workflow for BIOM, Newick plus sidecar abundance tables, and QIIME-exported files. MetaTree constructs a shared reference tree from the input hierarchy and uses it as a fixed node set, preserving one-to-one node correspondence so that samples, groups, and contrasts are compared on the same topology across views. Quantitative values are encoded using three visual channels: node size, node color, and branch width. Users explore the data through four analysis modes: Sample View, Group View, Group Comparison, and Comparison Matrix. Interactive controls support synchronized zooming, adaptive label culling, filtering, and branch collapsing or expansion. The MetaTree home page is shown in Fig. 2. MetaTree provides extensive, user configurable controls for figure customization, including visual encoding, layout, label density, and filtering thresholds. It supports export of publication-ready figures and the associated statistics tables, and it can be accessed online or run locally without installation.

**Figure 1.**
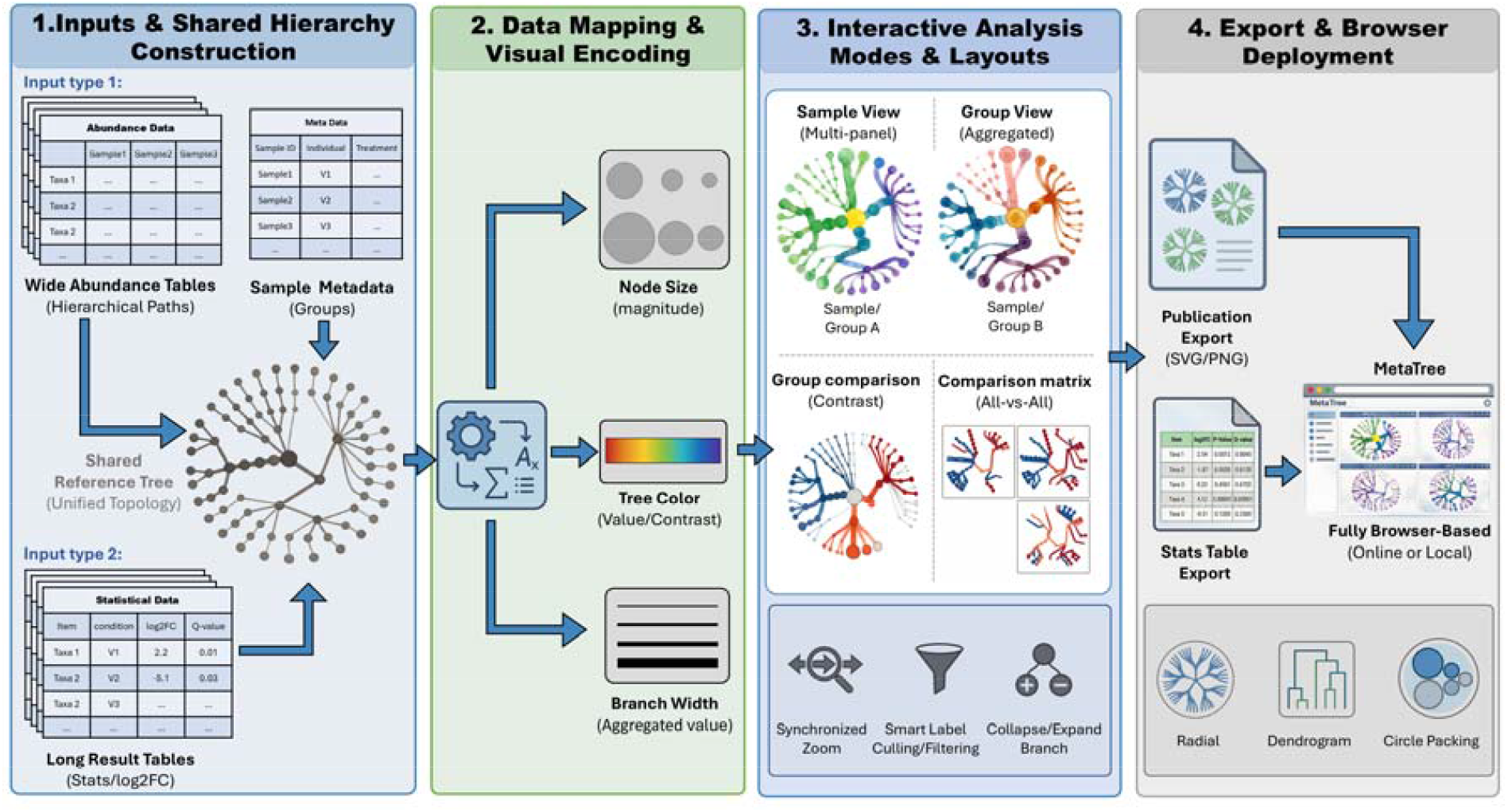
MetaTree interface overview and end-to-end workflow for hierarchical visualization and comparison. MetaTree visualizes hierarchical profiles on a shared reference topology for topology-consistent comparison. Node size/color and branch width encode abundance or contrasts.

**Figure 2.**
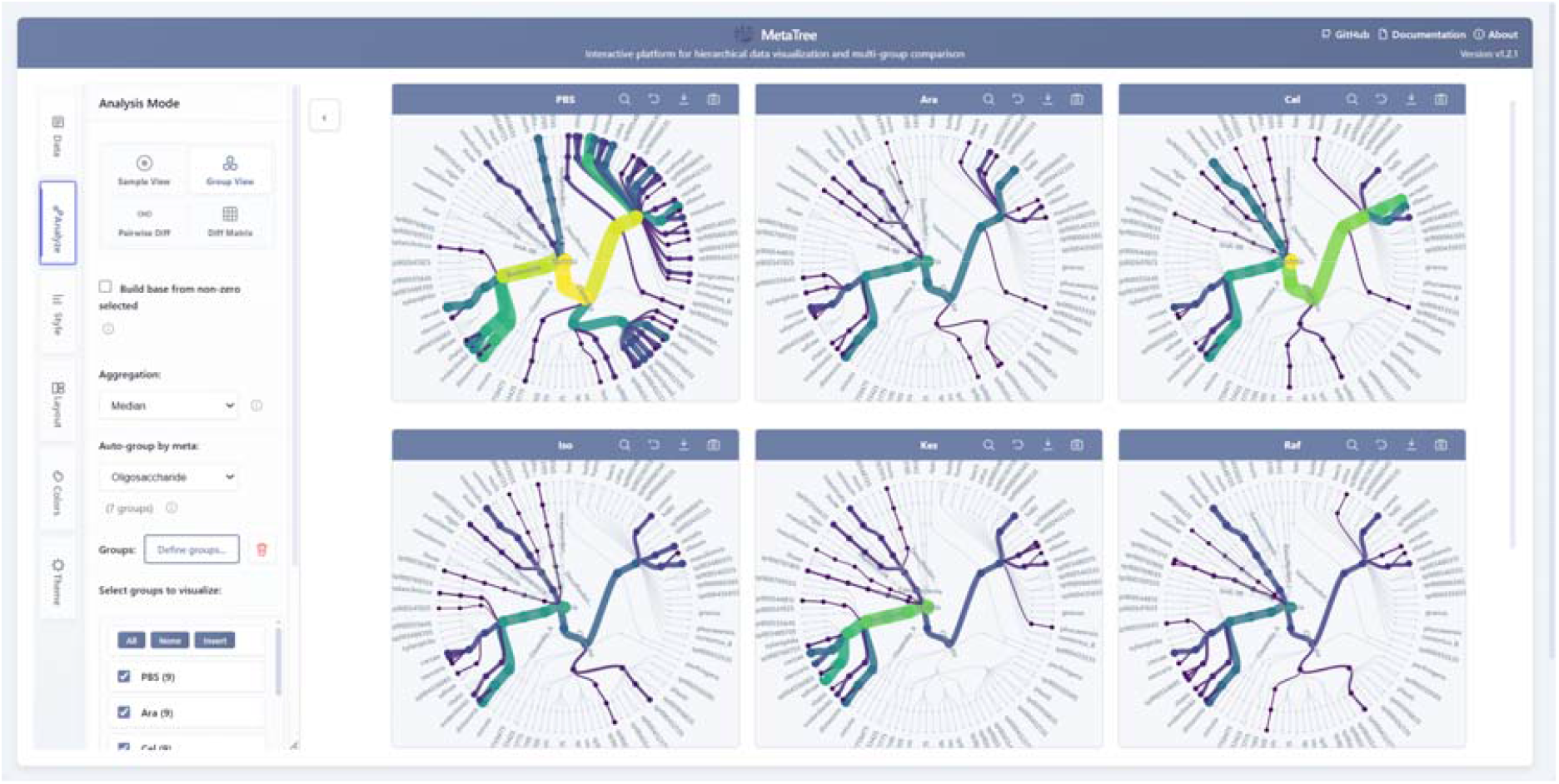
MetaTree graphical user interface (GUI). The example panels illustrate the visual grammar used throughout MetaTree: node color encodes the selected quantitative variable, node size reflects its magnitude, and branch width summarizes aggregate hierarchical signal. The GUI provides controls for data loading, grouping, visual encoding, filtering, synchronized navigation, and figure export.

### Exploration and comparison on a shared reference tree

To demonstrate MetaTree’s capabilities, we applied it to a previously published human gut microbiome metaproteomics dataset generated from stool-derived communities of three donors cultured in vitro with several structurally distinct oligosaccharides [27]. Taxonomic and operational taxon–function (OTF) intensity tables were generated using MetaX, a peptide-centric metaproteomics workflow [21], and used as input to MetaTree. In this dataset, donor identity is denoted as V1–V3, and we use these donor labels as example groups for illustrating group-level summaries and contrasts.

MetaTree provides four analysis modes on a shared reference tree, allowing hierarchical profiles and contrasts to be interpreted on a consistent topology: Sample View and Group View for profile exploration, Group comparison for two-group contrasts, and Comparison matrix for systematic multi-group screening. Sample View renders one heat-tree per sample as a panel set, whereas Group View aggregates replicates within user-defined groups and renders one heat-tree per group. Figure 3 highlights the comparison-oriented modes. Group View summarizes group-level patterns for V1 and V2 on the same topology with a shared color scale (Fig. 3a). Group comparison visualizes a selected contrast between two groups and can restrict the display to nodes passing user-defined significant thresholds (Fig. 3b). In this mode, MetaTree provides built-in per-node hypothesis testing using either a Wilcoxon rank-sum test or Welch’s t-test and applies BH correction to control FDR. For studies with more than two groups, Comparison matrix computes all pairwise contrasts and arranges them as a matrix of differential heat-trees for rapid screening; each cell can be opened for focused inspection (Fig. 3c).

**Figure 3.**
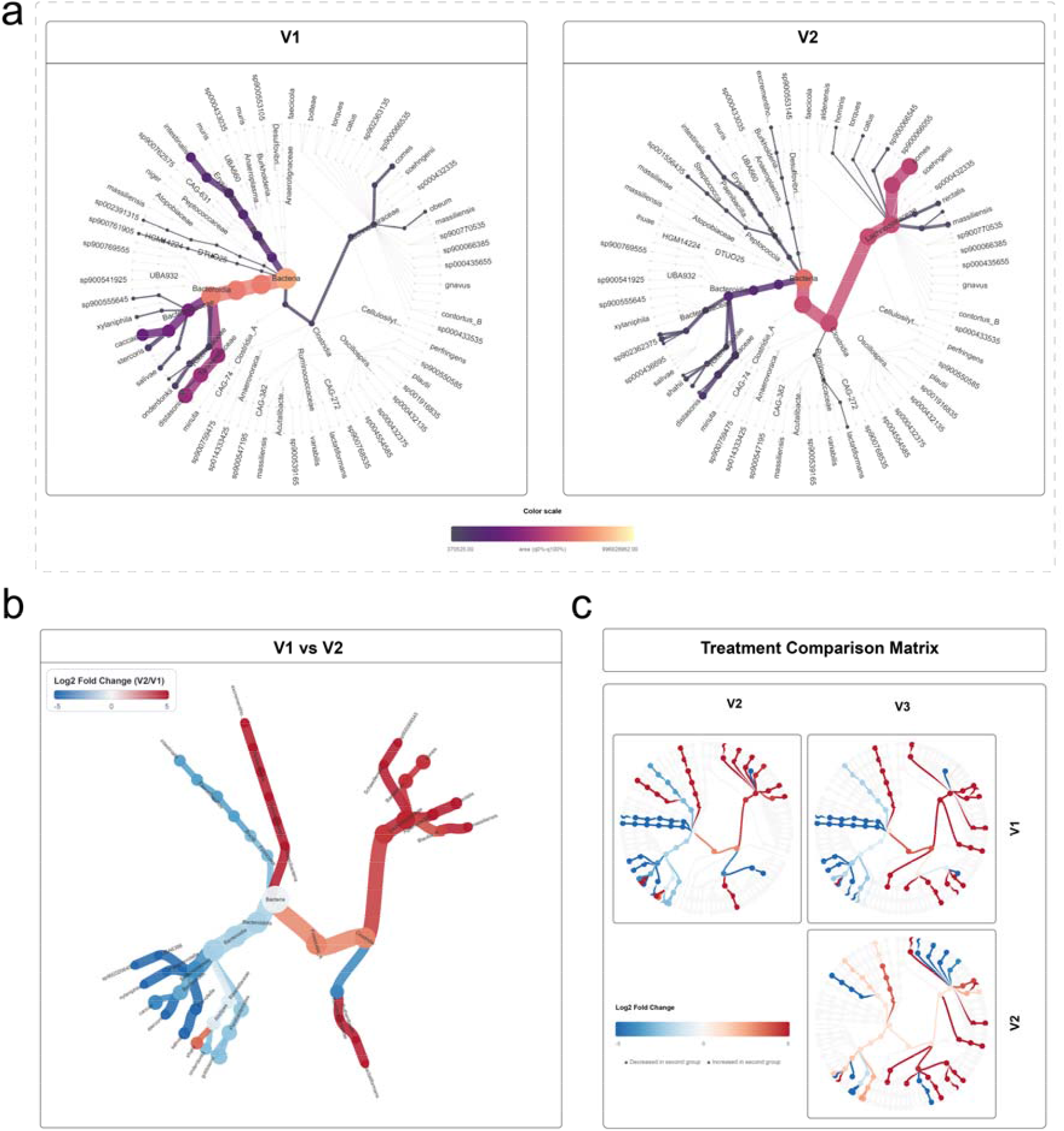
Group-level summaries and differential comparisons in MetaTree. **(a)** Group View: aggregated profiles for V1 and V2 on a shared topology; node color and node size encode group-level summarized intensity, and branch width reflects aggregate signal within each clade. **(b)** Group comparison: V1 versus V2 contrast; node color encodes log2 fold-change, while displayed nodes can be filtered by FDR and absolute log2 fold-change thresholds. **(c)** Comparison matrix: all pairwise contrasts among V1–V3, with each cell summarizing one pairwise group contrast on the same reference topology.

### Interactive controls for comparative visualization and multi-layout rendering

MetaTree provides GUI panels to configure data input, visual encoding, and figure layout (Fig. 4a). Hierarchical abundance tables and optional sample metadata can be loaded with automatic input parsing, including detection of common table delimiters and hierarchy separators, reducing manual preprocessing. Users can filter samples by metadata and restrict displayed features by name to focus on specific conditions or clades.

**Figure 4.**
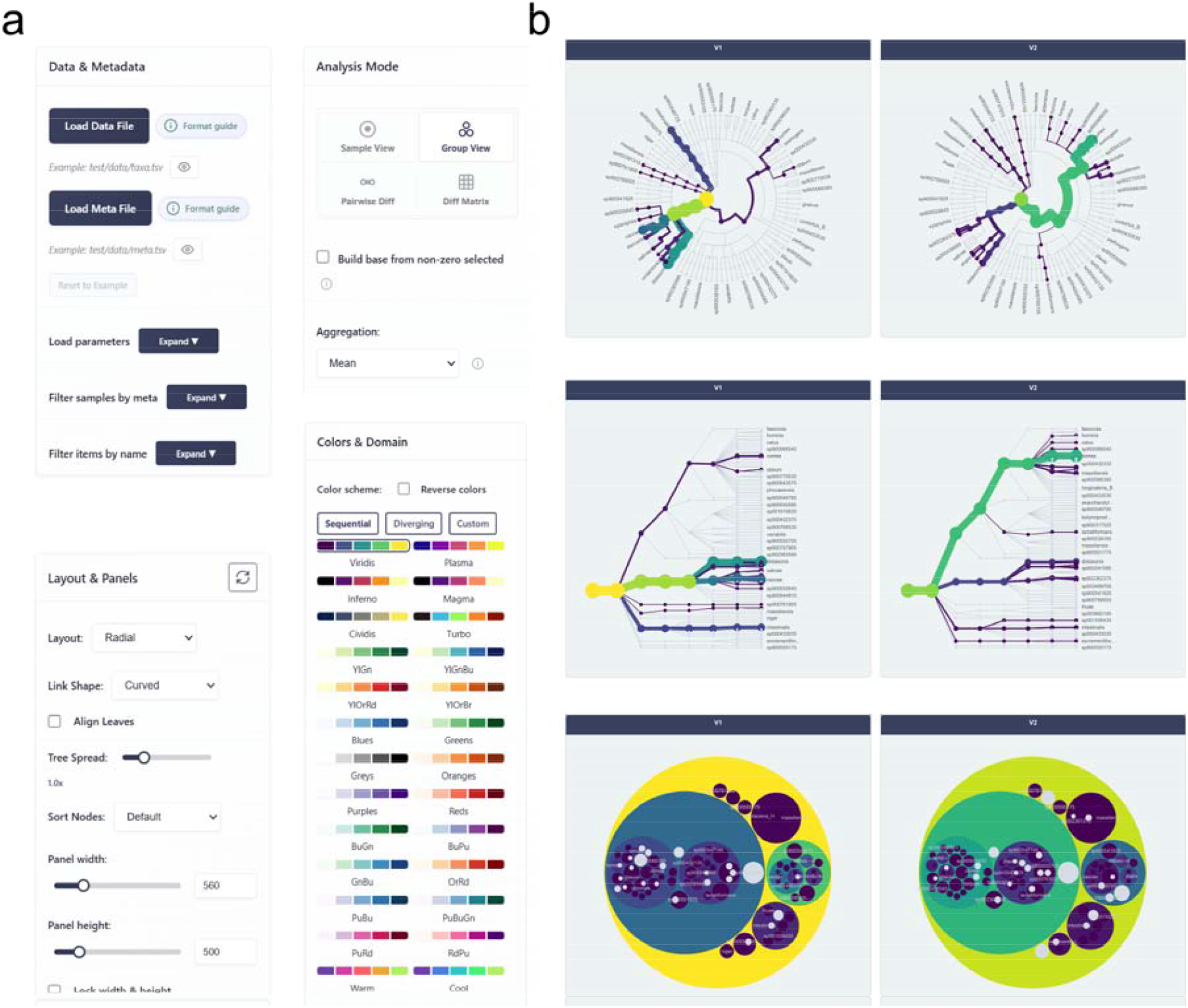
Interactive configuration panels and multi-layout rendering in MetaTree. **(a)** Control panels for data loading, filtering, label settings, and layout configuration. **(b)** The same dataset rendered with different layouts (radial tree, dendrogram tree, and circle packing).

Visual encoding is configurable via sequential and diverging palettes, optional palette reversal, and user-defined color domains to enforce consistent scaling across panels. Label presentation can be tuned through font and length limits, hierarchy-level selection, and automatic overlap culling for dense trees. In addition, label colors can be assigned automatically or customized to match user-defined groupings, improving interpretability without changing the underlying hierarchy.

MetaTree supports multiple layouts for the same hierarchy, including radial trees, dendrogram trees, and circle packing (Fig. 4b). Layout parameters and panel dimensions can be adjusted interactively. For comparative exploration, MetaTree enables coordinated multi-panel interaction with synchronized navigation (linked zoom, pan), and hover tooltips provide node-level details without adding persistent visual clutter.

### Performance and scalability benchmarking

To provide an objective assessment of browser-side performance, we benchmarked MetaTree across increasing hierarchy sizes and workloads in its four analysis modes using both dendrogram and radial layouts (Fig. 5). In Sample view and Group view, 10,000-node single-panel displays remained below 0.4 s, whereas 9-panel displays required approximately 2.2-2.7 s. Group comparison was less demanding, with 10,000-node initial render times of approximately 0.5-0.6 s, while Comparison matrix performance depended strongly on the number of displayed contrasts. Post-render JavaScript heap usage increased with hierarchy size and workload, reaching 85-104 MB in the largest 9-panel Sample view and Group view conditions. Full benchmark values are provided in Supplementary Table S1.

**Figure 5.**
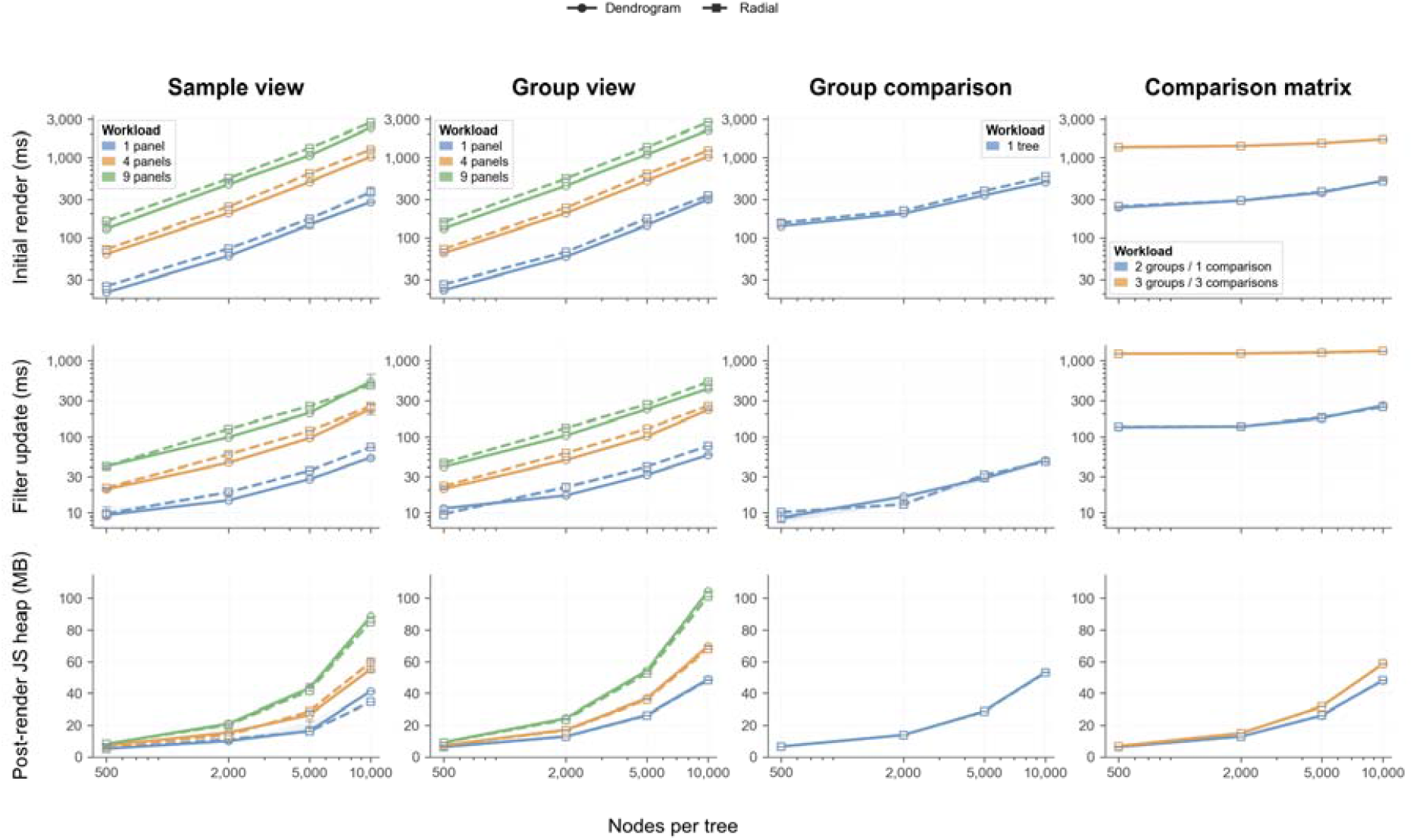
Performance and scalability benchmark of MetaTree across four analysis modes. Columns correspond to Sample view, Group view, Group comparison, and Comparison matrix. Rows show initial render time, filter-update time, and post-render JavaScript heap usage after initial rendering. Benchmarks were evaluated on hierarchies of approximately 500, 2,000, 5,000, and 10,000 nodes, with increasing workloads where applicable. Solid lines with circles indicate the dendrogram layout, and dashed lines with squares indicate the radial layout. Points represent medians across five repeated runs; error bars show interquartile ranges. Timing axes are shown on a logarithmic scale.

## Discussion

MetaTree is designed for systematic comparison and reporting of hierarchical quantitative profiles through a browser-based interface optimized for multi-panel inspection and figure production. Its central contribution is not simply tree rendering, but the use of a shared reference hierarchy that preserves one-to-one node correspondence across samples, groups, and contrasts. This design allows the same clade or category to be interpreted in the same topological position across views, thereby supporting more consistent comparison of hierarchical patterns across conditions. In addition to interactive exploration, MetaTree provides exportable figures and statistics tables, and can also be embedded into external workflows through lightweight programmatic integration.

As summarized in Table 1, our comparison was not intended as a systematic review of all microbiome visualization software. Instead, iTOL, Krona, and Metacoder were selected as representative baselines because they correspond to three conceptually distinct and practically relevant classes of hierarchical visualization workflow: annotation-centric tree presentation, interactive drill-down exploration of hierarchical composition, and heat-tree generation in a scripting environment. MetaTree is therefore best viewed as complementary to existing hierarchical visualization tools rather than as a replacement for all workflows. iTOL [15] is strong for annotation-rich presentation of user-supplied trees, and Krona [14] is effective for interactive drill-down of hierarchical composition. However, neither provides a native workflow for topology-consistent multi-panel comparison in which samples, groups, and contrasts are displayed on a shared reference hierarchy with synchronized navigation and direct all-versus-all screening. Metacoder [9] supports hierarchy-aware group comparison through heat-tree generation in an R-based environment, but it is primarily code-driven and oriented toward static figure generation; for large or dense hierarchies, maintaining label readability and assembling multi-panel comparative figures may require substantial manual tuning. MetaTree addresses these limitations through browser-based coordinated rendering across multiple layouts, adaptive label management, branch collapsing, linked navigation, and an integrated comparison matrix, while also supporting direct export of publication-ready figures and associated statistics tables.

**Table 1.** Comparison of MetaTree with representative hierarchical visualization tools.

| Capability | MetaTree | iTOL | Krona | Metacoder |
| --- | --- | --- | --- | --- |
| Topology-consistent comparison across groups | ? | — | — | ? |
| Coordinated multi-panel interaction | ? | — | — | — |
| No installation (web-based) | ? | ? | Δ | — |
| Multiple layouts | ? | ? | — | — |
| Interactive visual encoding controls | ? | ? | ? | — |
| Built in two group testing | ? | — | — | — |
| Pairwise comparison matrix (all-versus-all) | ? | — | — | — |
| Publication-ready export (vector and raster) | ? | ? | Δ | ? |

? supported natively; Δ possible with custom scripting or non-native workflows; — not supported.

MetaTree also provides built-in two-group statistical summaries for rapid exploratory screening. Users can project log2 fold-changes and adjusted P-values onto the hierarchy, apply user-defined thresholds, and export the resulting views and tables. This functionality is intended to support rapid identification of clades or categories that merit closer inspection. For confirmatory inference in complex experimental designs, study-appropriate statistical models can be fitted externally and the resulting statistics can then be imported into MetaTree for hierarchical interpretation and reporting. Although the main examples in this study are biological, the MetaTree framework is not restricted to biological hierarchies. It can in principle be applied to rooted hierarchies in which quantitative values are associated with hierarchical items or nodes. To illustrate this generality, we applied MetaTree to a non-biological economic dataset derived from World Bank GDP data, in which countries were organized within a rooted world-region hierarchy and visualized across two time points (Fig. 6). This example shows that the same topology-consistent comparison framework can also be used for hierarchical quantitative datasets outside microbiome or metaproteomics applications.

**Figure 6.**
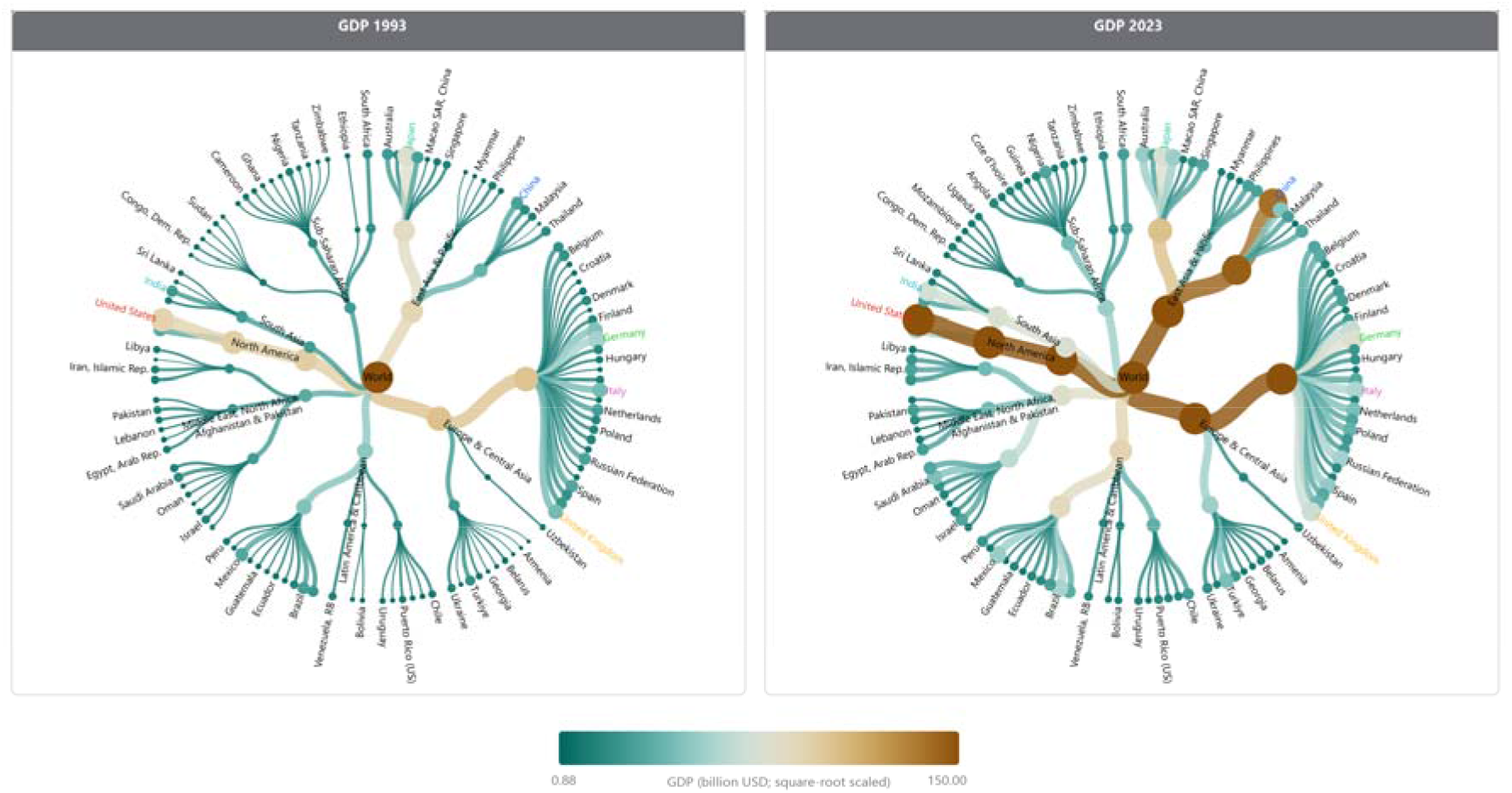
Illustrative use of MetaTree for a non-biological hierarchical economic dataset. Countries were arranged within a rooted world-region hierarchy and visualized using World Bank GDP values for 1993 and 2023. This example was included to illustrate that MetaTree is not restricted to biological data and can also be used for other rooted hierarchical quantitative datasets.

MetaTree is intended as a lightweight, installation-free platform for interactive exploration and reporting rather than as a confirmatory statistical framework. Its built-in two-group comparisons should therefore be interpreted as exploratory. Because tests are performed at the node level on a shared hierarchy, including internal nodes whose values are derived from their descendants, parent and child nodes are structurally dependent and the resulting tests are not independent. The BH procedure is used as a practical default for interactive filtering, but it does not explicitly account for hierarchy-induced dependence. More formal tree-structured multiple-testing strategies, such as TreeFDR, may provide a more principled way to address this issue. At the same time, MetaTree is not restricted to its built-in statistical workflow, because externally computed statistical results can also be imported and interpreted within the same hierarchical framework. This allows users to apply study-specific models or more rigorous statistical procedures outside MetaTree and then use MetaTree for topology-aware visualization and reporting. In addition, interpretation depends on the quality and appropriateness of the input hierarchy, and extremely dense trees may still challenge readability despite label culling, filtering, and branch collapsing. Benchmark results nevertheless indicate that browser-side performance scales predictably across the tested hierarchy sizes and workloads, supporting practical use for interactive exploration. Future work will focus on improving reproducibility through provenance capture for exported figures, extending comparison options and format compatibility, and further exploring statistically rigorous methods that account for hierarchical dependence.

## Conclusion

MetaTree is an installation-free web platform for topology-consistent visualization, exploration, and comparison of hierarchical quantitative profiles. It supports coordinated multi-panel exploration, automated comparison matrices, and integration of statistical results with interactive filtering and thresholding. These capabilities enable rapid generation of publication-ready figures for microbiome and other hierarchical datasets.

## Availability and requirements

Project name: MetaTree

Project home page: https://github.com/byemaxx/MetaTree

Operating system(s): Platform independent

Programming language: JavaScript

Other requirements: A modern web browser

License: MIT

Any restrictions to use by non-academics: No

## List of abbreviations

BH: Benjamini-Hochberg
D3.js: Data-Driven Documents (JavaScript library)
FDR: false discovery rate
GUI: graphical user interface
HTML: HyperText Markup Language
iTOL: Interactive Tree Of Life
log2FC: log2 fold change
OTF: operational taxon–function
PNG: Portable Network Graphics
rRNA: ribosomal RNA
SVG: Scalable Vector Graphics

## Declarations

### Ethics approval and consent to participate

Not applicable.

### Consent for publication

Not applicable.

### Availability of data and materials

MetaTree is available as a hosted web application (https://byemaxx.github.io/MetaTree). Its source code, benchmark workflows and outputs, documentation, and example files, together with instructions for optional local deployment, are available on GitHub (https://github.com/byemaxx/MetaTree). The example dataset used in this study was derived from the previously published human gut microbiome metaproteomics study by Zhang et al. [27], and the underlying mass spectrometry proteomics data are publicly available in PRIDE under dataset accession PXD064733.

### Authors’ contributions

Q.W. designed the method, developed the software, and drafted the manuscript. A.Z. provided key insights, tested the software, and reviewed the manuscript. Z.N. contributed to methodological enhancements and reviewed the manuscript. D.F. reviewed and provided feedback on the manuscript.

## Acknowledgements

We used a large language model (ChatGPT, OpenAI) to assist with English language editing for clarity and readability. All scientific content, analyses, and conclusions were produced and verified by the authors.

